# Improved mouse models of the small intestine microbiota using region-specific sampling from humans

**DOI:** 10.1101/2024.04.24.590999

**Authors:** Rebecca N. Culver, Sean Paul Spencer, Arvie Violette, Evelyn Giselle Lemus Silva, Tadashi Takeuchi, Ceena Nafarzadegan, Steven K. Higginbottom, Dari Shalon, Justin Sonnenburg, Kerwyn Casey Huang

## Abstract

Our understanding of region-specific microbial function within the gut is limited due to reliance on stool. Using a recently developed capsule device, we exploit regional sampling from the human intestines to develop models for interrogating small intestine (SI) microbiota composition and function. *In vitro* culturing of human intestinal contents produced stable, representative communities that robustly colonize the SI of germ-free mice. During mouse colonization, the combination of SI and stool microbes altered gut microbiota composition, functional capacity, and response to diet, resulting in increased diversity and reproducibility of SI colonization relative to stool microbes alone. Using a diverse strain library representative of the human SI microbiota, we constructed defined communities with taxa that largely exhibited the expected regional preferences. Response to a fiber-deficient diet was region-specific and reflected strain-specific fiber-processing and host mucus-degrading capabilities, suggesting that dietary fiber is critical for maintaining SI microbiota homeostasis. These tools should advance mechanistic modeling of the human SI microbiota and its role in disease and dietary responses.

## Introduction

Humans depend on the dense and diverse community of microbes along the entire length of the intestinal tract, collectively known as the gut microbiota, for food digestion, immune system regulation, and protection against pathogens, among other important functions^1^. An important, yet often overlooked, aspect of the gut microbiota is regional heterogeneity and its impact on local physiology^2^. Driven by gradients of nutrient availability and other environmental factors, the microbial communities in each intestinal region are distinguished by specialized functions, metabolomes, immune niches, and proteomes^3–5^. The small intestine (SI) and its microbiota are linked to development of many diseases, including gastrointestinal cancer, metabolic syndrome, inflammatory bowel disease, and Celiac disease^6^. However, the human SI microbiota is not well characterized due to our reliance on analyzing feces, which is readily available but poorly represents microbial communities in more proximal gut regions^3,7,8^, and on conventional mouse microbiotas^9^, which may not accurately reflect the dynamics of the human microbiota. Thus, deeper understanding of the human SI microbiota and the features that distinguish it from the stool microbiota will require development and characterization of tools to directly probe and modify the SI microbiota and its impact on the host.

Current data regarding the human SI microbiota are typically from terminal sampling^7^ and lack information at the strain level that may be important for function. Human ileal stoma microbiotas showed that sub-strain proportions fluctuate over time, and highlighted diet as a key factor shaping microbiota composition in the small intestine^10^. In conventional mice, strain-specific features of SI microbes regulate dietary fat digestion and absorption by the host^11^. The pathobiont *Enterococcus gallinarum* exhibited site-specific within-host evolution, adapting to evade detection and clearance by the immune system and to enable liver translocation^12^. These examples highlight the importance of SI-specific host and microbe functions and motivate a systematic analysis of the regional colonization preferences of human-associated microbes.

Although humanized mice (ex-germ-free animals colonized with consortia of human gut commensals) have proven a powerful tool for probing human stool microbiota dynamics and physiology, the extent to which the SI microbiota is accurately modeled remains to be explored due to limited access to human SI microbiota samples. Humanization of germ-free mice requires sufficient material for colonizing multiple animals per experiment; while stool is readily available, stable, and abundant, the human SI is a highly variable environment that is difficult to sample in large volumes. Stool-derived *in vitro* communities have proven stable and reproducible over repeated passaging^13^, able to predict the ability to metabolize drugs^14^, and representative of the community composition, host proteome, and microbiota response to antibiotics in humanized mice^13^. Such “top-down” communities are replenishable via *in vitro* propagation and hence can remedy issues with inoculation supply. Furthermore, highly diverse, synthetic (defined) communities of stool isolates stably colonize gnotobiotic mice and can be used *in vitro* and *in vivo* to study microbiota functions such as colonization resistance^15^ and the effects of diet^16^, and are precursors to microbial therapeutic cocktails. However, previous studies of synthetic communities have largely involved stool isolates. Therefore, the impact of microbes that are resident in the SI and their site-specific effects within the intestinal tract remains to be explored.

We recently developed and clinically evaluated an ingestible device that collects samples along the human intestinal tract during normal digestion^8,17^. Using >240 intestinal samples from 15 healthy subjects, we identified significant differences in bacterial composition and gene content, prophage induction, host proteins, and metabolites such as bile acid profiles between the intestines and stool^8,17^.

Moreover, we found that dietary metabolites strongly segregated between the upper and lower intestinal tract^17^. Previous research has focused on the relationship between microbially accessible carbohydrates (MACs), a component of dietary fiber and driver of gut microbial metabolism, and the colonic/stool microbiota^18,19^. Fiber deprivation increases the abundance of mucus-degrading bacteria in the gut, leading to increased pathogen susceptibility^16^. Mucus structure and thickness varies with diet^20^, and differs dramatically between the SI and colon^21^. This physiological and nutrient diversity along the intestinal tract has the potential to drive large ecological variations, but the role of diet on the human SI microbiota remains largely unexplored. Bacteria collected from the SI by our ingestible devices largely remained viable^8^, indicating that these samples are amenable to cultivation of *in vitro* communities and isolation of phylogenetically diverse bacterial species for study of SI colonization and links to diet in gnotobiotic mice.

Here, we show that *in vitro* culturing of human SI contents results in stable, reproducible, representative communities. Germ-free mice colonized with *in vitro* communities derived from human SI versus stool samples exhibited distinct gut microbiota compositions. The reproducibility of SI microbiota composition across mouse experiments increased dramatically when mice were colonized with a human SI-derived *in vitro* community. Switching from a standard to a MAC-deficient diet led to differential restructuring of the stool and SI microbiota. To empower interrogation of these colonization differences, we assembled a library of >500 strains that collectively capture the diversity of the SI microbiota, spanning 19 families and all major phyla of the gut microbiome of a single subject. Colonization with synthetic communities revealed that taxa maintain preference for the spatial niche from which they were isolated. Functional potential varied with diet, intestinal location, and colonization source, highlighting the importance of these communities and isolates as models of the human SI microbiota.

## Results

### *In vitro* culturing of human intestinal contents generates stable, reproducible, representative communities

To elucidate the physiology and functions of the microbial communities throughout the human intestinal tract, we capitalized on the observation that most bacterial cells sampled by our ingestible devices remain viable^8^. To determine whether *in vitro* passaging of unfractionated SI contents would generate diverse *in vitro* communities suitable for modeling the human SI microbiota, we acquired samples from three regions of the intestinal tract of a healthy human subject as well as a matching stool sample (four samples total, Fig. 1A). After the devices were collected, they were transported within one hour to an anaerobic chamber along with the stool sample, and their contents were inoculated into pre-reduced Brain Heart Infusion (BHI) medium (Fig. 1A; Methods). In a previous study, we determined that BHI best recapitulated the composition and diversity of a wide range of inoculating stool samples among a panel of commonly used media for gut commensal culturing^13^. Motivated by the known variability in microbiota composition and pH along the human SI^22,7,8^, we passaged all samples across a wide range of physiologically relevant pH values to identify conditions that improve modeling of the SI microbiota. The pH of the inoculating intestinal samples ranged from 5.0-7.1; we varied the initial pH of the medium from 4 to 10. At pH 5, communities were dominated by a single *Lactobacillus* species and were therefore ignored hereafter due to their lack of diversity (Fig. S1A); at pH 4, essentially no growth was observed (data not shown).

**Figure 1:**
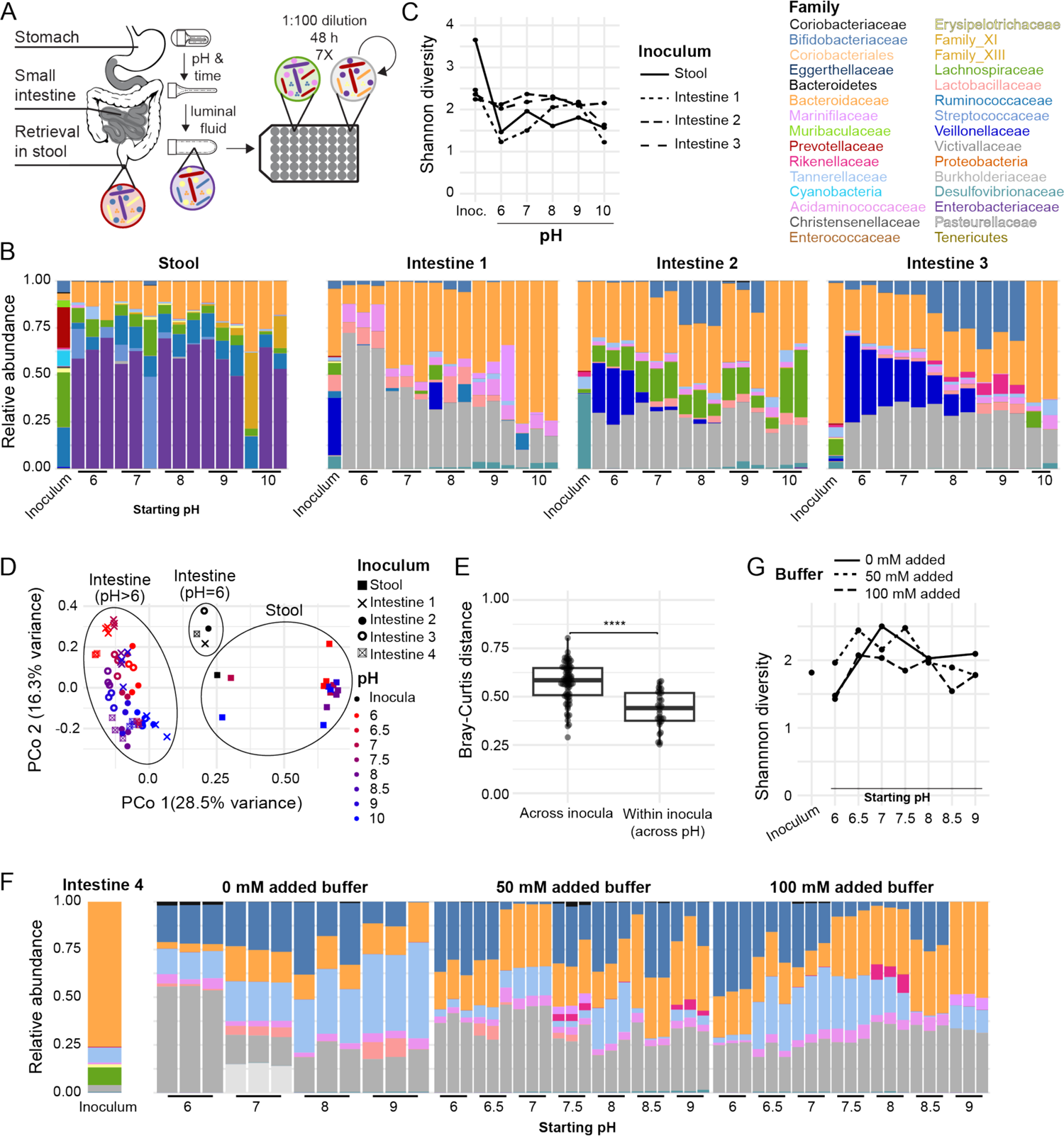
*In vitro* communities derived from human small intestinal samples are diverse and vary with both inoculum and pH. A) Schematic of *in vitro* community culturing. Three noninvasive sampling devices that collect luminal fluid from various locations in the intestinal tract were retrieved from stool, and immediately transported to an anaerobic chamber for inoculation into 96-well plates. Cultures were passaged through multiple cycles of dilution and growth to ensure stability, with various starting pH values and buffer concentrations. B) *In vitro* community composition depends on both the inoculum and starting pH of the medium. Shown are family-level relative abundances for the inocula and three replicate communities for each pH. C) The average Shannon diversity across replicates (*n*=3) of the *in vitro* communities varied somewhat across starting pH values, and communities derived from SI samples, but not stool, maintained the diversity of the inoculum at certain pHs. D) Principal coordinate analysis based on Bray-Curtis distance among *in vitro* community and inoculum compositions. Colors represent the starting pH of the medium for *in vitro* communities, and the initial inocula are black. Symbol denotes the inoculum. E) Bray-Curtis distance between the composition of pairs of *in vitro* communities derived from distinct inocula varied significantly more than that of *in vitro* communities derived from the same inoculum at different starting pH values (****: *p*≤0.0001, two-sided Wilcoxon rank-sum test). F) Family-level relative abundances for a separate small intestinal inoculum from the same subject as in (B-E) and the *in vitro* communities derived from this sample at various starting pH values and buffer concentrations (*n*=3 replicates). G) The average Shannon diversity across replicates (*n*=3) of the *in vitro* communities derived from the SI sample used in (F) was similar to that of the inoculum but varied across starting pH values and buffer concentrations.

We used 16S rRNA gene sequencing to assess the relative abundance of each taxon in the inocula and *in vitro* communities (Fig. 1B). Representatives of 17 families were reliably detected (>10^-3^ abundance) in the stool inoculum and 9-18 families were detected in the three SI inocula. Yield, as assessed by optical density, varied by <2-fold across all *in vitro* communities, indicating that relative abundances could be at least approximately compared across communities^23^.

Community composition equilibrated quickly (Fig. S1B), and technical replicates reached highly similar compositions (Fig. S1C). In the *in vitro* community derived from the stool sample, the initial pH of the medium had little effect on community composition at the family level. By contrast, in certain SI-derived *in vitro* communities, increasing the pH of the medium at the beginning of each passage led to family-specific abundance changes, such as a decrease in Veillonellaceae and increases in Bifidobacteriaceae and Rikenellaceae (Fig. 1B). The Shannon diversity of the communities varied substantially, with some communities exhibiting diversity almost as high as the inoculum (Fig. 1C). Overall, communities were robust to pH variations in the intermediate ranges.

Communities derived from the intestinal samples retained representatives of 7-11 families constituting 78%-99.6% of the family-level abundance from the corresponding inoculum, indicating that *in vitro* communities can preserve a large fraction of the SI microbiota. In a Principal Coordinates Analysis (PCoA), SI-derived communities clustered away from stool-derived communities (Fig. 1D). Clustering of the *in vitro* communities was driven more by inoculum than by pH during passaging (Fig. 1D,E), indicating that *in vitro* culturing of a microbial community intended to be representative of a particular location in the human intestinal tract requires collection from the desired site. As SI samples are not as dense nor as large in volume as stool samples, culturing microbial communities from the SI is a useful means for obtaining sufficient biomass for scientific study and therapeutic applications.

Since bacterial modification of environmental pH could affect community growth dynamics, we derived *in vitro* communities with 0 mM, 50 mM, or 100 mM of a buffer effective at each starting pH and investigated initial medium pH values from 6-9 in increments of 0.5. Given the limitations in device sample volume, we acquired another sample from the upper SI of the same subject to carry out these experiments (Methods). As intended, buffer addition restricted the ability of the microbes to modify environmental pH (Fig. S2). At the family level, several media conditions produced communities with composition and diversity resembling the initial inoculum (Fig. 1F,G). The diversity of communities derived at pH<8 with 50 mM buffer was generally higher compared to without buffer due to the appearance of several families that were either not detected or were at low abundance in the initial inoculum (Fig. 1F,G); for instance, in the community derived at pH 7.5 + 50 mM buffer, Coriobacteriaceae and Parasutterellaceae emerged from below the limit of detection and Acidaminococcaceae and Rikenellaceae increased several fold in relative abundance. The addition of 100 mM buffer largely prevented the emergence of families that were not detected in the inoculum, suggesting that pH modulation by other community members was important for their growth. Thus, medium buffering can affect *in vitro* culturing and increase the similarity between an *in vitro* community and its initial inoculum.

### Intestinal microbiota composition in gnotobiotic mice is dependent on the origin of the colonizing community

To develop an improved mouse model of SI colonization by human microbiota, we first sought to quantify colonization differences between communities derived from the SI and stool of the same human subject. We utilized two of the *in vitro* communities described above: an SI-derived community passaged in BHI at pH 7.5 + 100 mM buffer (henceforth referred to as SICom) and a stool-derived community passaged in BHI at pH 7 (henceforth, StoolCom) (Fig. 2A). The intermediate pH was selected to avoid inhibition of the growth of taxa sensitive to acidic environments, and the 100 mM buffer concentration was selected to determine if suppression of low abundance members *in vitro* would exclude them from mouse colonization as opposed to microbes below the level of detection *in vitro* blooming in the context of the mouse GI tract. SICom contained 6 of the 10 families detected in the SI inoculum, collectively representing 87.9% of the family-level abundance, and its family-level composition was correlated with that of the SI inoculum from which it was derived (*R*=0.69, *p*=0.003, Pearson correlation). StoolCom contained 5 of the families detected in the stool inoculum that collectively represent 54.4% of the family-level abundance. Taken together, these communities provide the potential for a diverse range of colonization outcomes to study regional specificity along the intestinal tract.

**Figure 2:**
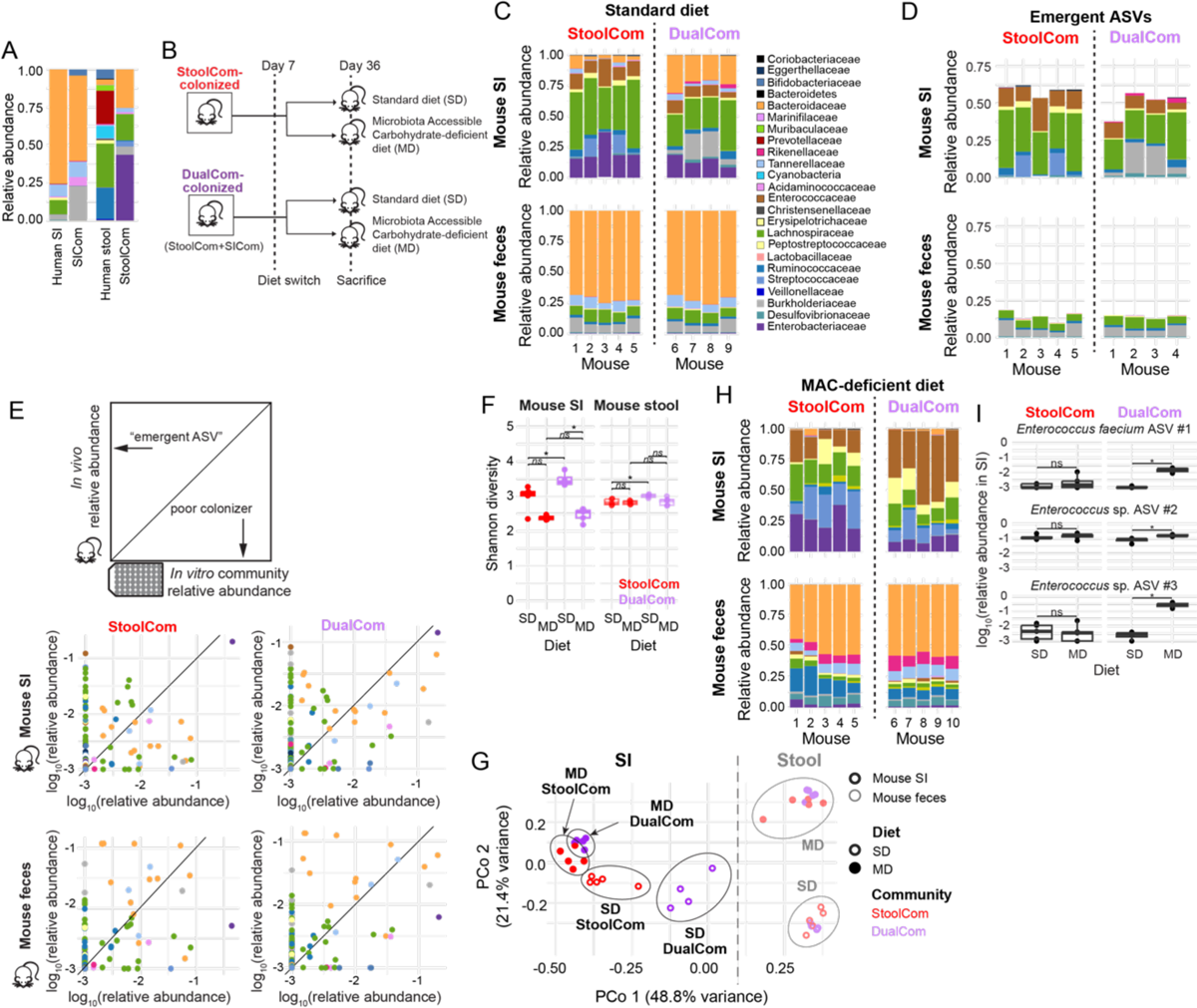
Spatially distinct *in vitro* communities create distinct colonization states along the intestinal tract of mice fed either a standard or MAC-deficient diet. A) Family-level relative abundances for the initial human intestinal and stool sample with corresponding *in vitro* communities derived from either the human intestine (SICom) or human stool (StoolCom). Many, though not all, of the prominent families were preserved in the *in vitro* communities. B) Schematic of mouse experiment in which two isolators each contained two cages of five mice each. In the first isolator, both cages were gavaged with StoolCom, and after seven days, one of the cages was switched from a standard diet (SD) to a MAC-deficient diet (MD). The same process was followed in the second isolator, but with DualCom (an equal volume mixture of SICom and StoolCom). After 36 days, all mice were sacrificed and feces and intestinal contents were collected. C) Family-level relative abundance was highly distinct between the mouse SI (top) and fecal microbiotas (bottom) for StoolCom-(left) and DualCom-colonized (right) SD-fed mice. Each bar represents an individual mouse. D) The abundance of emergent ASVs was higher in the mouse SI microbiota compared to the fecal microbiota regardless of colonization condition. Family-level relative abundance of emergent ASVs detected in the mouse SI (top) and fecal microbiotas (bottom) for StoolCom-colonized (left) and DualCom-colonized (right) SD-fed mice. Each bar represents an individual mouse. E) Comparison between *in vitro* community inoculum (SICom (left) or DualCom (right)) and mouse SI (top) or fecal (bottom) microbiota composition of SD-fed mice reveals both emergent and robustly colonizing ASVs. Each circle represents the mean log_10_(relative abundance) of an ASV across mice, and color corresponds to family-level annotation. Shown are Pearson correlation coefficients. F) DualCom-colonized mice had significantly higher Shannon diversity along the intestinal tract when fed a SD but not a MD. The Shannon diversity for each mouse intestinal microbiota across all colonization and diet conditions is shown. *p*-values are from a two-sided Wilcoxon rank sum test with Benjamini-Hochberg correction. *ns*: not significant; *: *p*≤0.1. G) Family-level relative abundance was highly distinct between the mouse SI (top) and fecal microbiotas (bottom) for StoolCom-(left) and DualCom-colonized (right) MD-fed mice. Each bar represents an individual mouse. H) Principal coordinate analysis based on Bray-Curtis distance between mouse microbiotas based on ASV relative abundance. Each light/dark circle represents a mouse fecal/SI microbiota, respectively. The color denotes the colonization condition, and the shape indicates the diet. I) The log_10_(relative abundance) of three *Enterococcus* ASVs in the SI was diet dependent in DualCom-colonized (left) but not StoolCom-colonized (right) mice. *p*-values are from a two-sided Wilcoxon rank sum test with Benjamini-Hochberg correction. *ns*: not significant; *: *p*≤0.1.

Since stool is likely to contain low levels of SI-derived bacteria, we sought to assess the ability of a stool-derived community alone to colonize the mouse SI as compared to a stool-derived community combined with an SI-derived community. We colonized cohorts of germ-free mice fed a standard diet (SD) with StoolCom only or SICom plus StoolCom (hereafter, DualCom) in separate isolators (Fig. 2B). After 36 days, fecal samples were collected, and the mice were sacrificed to access SI contents. The SI was divided into 3 sections (proximal, mid, and distal), and intestinal contents from each section were collected. 16S sequencing was used to quantify microbiota composition in each sample. Fecal microbiota composition on day 36 was highly correlated across mice within a cage and differed dramatically from SI microbiota composition (Fig. 2C). Although extensive spatial variability has been observed in human intestinal microbiota^7,8^, composition at the ASV level was largely correlated between the proximal, mid, and distal regions of the mouse intestine. For our analyses, reads from these three regions were summed to represent one intestinal sample per mouse.

We next assessed whether only taxa that were abundant in the inoculum could colonize the mouse intestinal tract. Of the eight families detected in the StoolCom inoculum, all eight were detected in StoolCom-colonized mice: six in the SI and seven in the feces. Unexpectedly, some ASVs undetectable in the StoolCom inoculum were detected in the SI and feces of the mice (hereafter referred to as “emergent”). On average, these emergent ASVs accounted for 58.9% of the bacterial abundance in the SI and 14.2% in the feces of StoolCom-colonized mice (Fig. 2D,E). Of the 10 families detected in the DualCom inoculum, seven were detected in the SI of DualCom-colonized mice and the same families were detected in the mouse feces. On average, emergent ASVs accounted for 50.2% of the bacterial abundance in the SI and 14.3% in the feces of DualCom-colonized mice (Fig. 2D,E). Interestingly, the taxonomic distribution of the emergent ASVs in the SI differed substantially between StoolCom- and DualCom-colonized mice (Fig. 2D). The high abundance of emergent ASVs in the mouse SI in both colonization conditions suggests that the SI environment applies selective pressures to the community distinct from *in vitro* culturing that are not apparent from analyses of feces, such that low abundance (undetectable) ASVs are still able to colonize the SI despite suppression of diversity through pH control *in vitro*.

The fecal microbiota composition of StoolCom-colonized mice was highly similar to that of DualCom-colonized mice (*R*=0.82, *p*<10^-13^, log_10_(ASV abundance), Pearson correlation; Fig. 2C), indicating that the addition of SI microbes to StoolCom had minimal impact on the distal gut. By contrast, the mouse SI microbiota composition was more weakly correlated (albeit still highly significant) between StoolCom- and DualCom-colonized mice (*R*=0.50, *p*<10^-13^, log_10_(ASV abundance), Pearson correlation; Fig. 2C). Both the SI and fecal microbiota of DualCom-colonized mice had significantly higher Shannon diversity than those of StoolCom-colonized mice (Fig. 2G), potentially reflecting inadequate niche filling using StoolCom alone. A PCoA further emphasized the similarity between stool microbiotas and the distinctiveness of SI microbiotas across colonization conditions (Fig. 2H). Thus, co-inoculation with SICom alters the composition of the mouse SI microbiota and increases diversity throughout the intestinal tract relative to StoolCom alone.

We previously identified five taxa enriched across the intestines of 15 human subjects: species from the *Escherichia/Shigella*, *Enterococcus*, *Bacteroides*, and *Romboutsia* genera^8^; *Romboutsia* species were not detected in any of the samples used to derive the *in vitro* communities. To evaluate whether the SI microbiota of these mice is a reasonable model of the SI microbiota in humans, we identified taxa that were significantly enriched (Methods) in the mouse SI in each colonization condition. StoolCom-colonized mice contained several ASVs that were significantly enriched in their SI compared to feces, including an *Escherichia/Shigella* and an *Enterococcus* species. An *Escherichia/Shigella* and *Enterococcus* species were also enriched in the SI of DualCom-colonized mice, as was a Desulfovibrionaceae species closely related to *Bilophila wadsworthia*, suggesting that DualCom produces more representative SI colonization than StoolCom alone. Taken together, colonization with communities derived from both human stool and the human SI leads to SI colonization of species that are generally enriched in that location.

### A fiber-free diet restructures the SI microbiota

Previous studies of the effects of a fiber-deficient diet in mice have highlighted restructuring of the SI and fecal microbiota^11,24^. However, these studies utilized specific pathogen free (SPF) mice with a conventional microbiota or gnotobiotic mice colonized with synthetic communities of isolates not necessarily representative of the human SI microbiota. Using our *in vitro* communities, we sought to interrogate how a MAC-deficient diet (MD) alters the SI microbiota relative to a SD. In separate isolators, we colonized two cages of mice with either StoolCom or DualCom and fed them a SD. Seven days later, one of the cages was switched to a MD (Fig. 2B). Feces was sampled periodically, and mouse SI contents were collected at the time of sacrifice (Fig. 2B). Similar to SD-fed mice, the SI microbiota composition of MD-fed mice was highly correlated along the intestinal tract, and all reads from MD-fed mouse SI samples were summed for each mouse into a single sample.

16S sequencing revealed numerous compositional changes after the diet switch in both the mouse SI and feces (Fig. 2G). The shift in the fecal microbiota after a diet switch was similar regardless of colonization state. Consistent with previous studies^16^, many fiber-degrading species decreased in relative abundance, including several *Bacteroides* species. Other species, including Lachnospiraceae members *Blautia massiliensis* and *Dorea longicatena, Anaerostipes hadrus*, *Sutterella wadsworthensis*, and *Phascolarctobacterium faecium*, also significantly decreased in the feces, suggesting that the lack of MACs or the resulting reduction in the levels of other metabolites such as short-chain fatty acids (SCFAs)^25^ influenced their abundance.

The SI microbiota of StoolCom- and DualCom-colonized mice were more similar to each other when fed a MD compared to a SD (Fig. 2C,G,H). Ten ASVs in the SI of StoolCom-colonized mice and 24 ASVs in the SI of DualCom-colonized mice were enriched by the diet switch, with an overlap of 9 ASVs. Many of the ASVs were enriched in SD-fed compared to MD-fed mice regardless of location (SI versus feces), including the fiber-degrading *B. ovatus* and SCFA-responsive *A. hadrus* and *Sutterella* species, suggesting that the switch between high-fiber and MAC-deficient diets affects the ecology of both the SI and distal colon.

In DualCom-colonized mice, the abundance in the SI of two *Enterococcus* ASVs was >7-fold enriched in MD-fed versus SD-fed mice (Fig. 2I). In StoolCom-colonized mice, both ASVs were detected in the SI but did not increase significantly in abundance after the switch to a MD (Fig. 2I). This contrast in behaviors suggests that SICom and StoolCom may contain distinct *Enterococcus* strains whose different functional capacities impact their response to dietary perturbations; we will return to this hypothesis later.

### Co-inoculation with SI microbes improves the reproducibility of mouse SI colonization

Given the high fraction of the mouse SI microbiota represented by emergent ASVs (Fig. 2D), we sought to test reproducibility of our observations across experiments. In a separate experiment, we introduced SICom and StoolCom to two cohorts of germ-free mice in separate isolators (Fig. 3A; henceforth, SICom-colonized and StoolCom^(2)^-colonized mice). The composition of each inoculum for this second set of experiments was highly correlated with the corresponding inoculum used for gavage in the first set of experiments in Fig. 2 (Fig. S3A,B), as expected based on the reproducibility of *in vitro* passaging^13^. After mouse sacrifice, feces and three regions of the SI were sampled for 16S sequencing. As before, the microbiota composition of the proximal, mid, and distal regions of the intestines were highly correlated with each other, and reads from these regions were summed.

**Figure 3:**
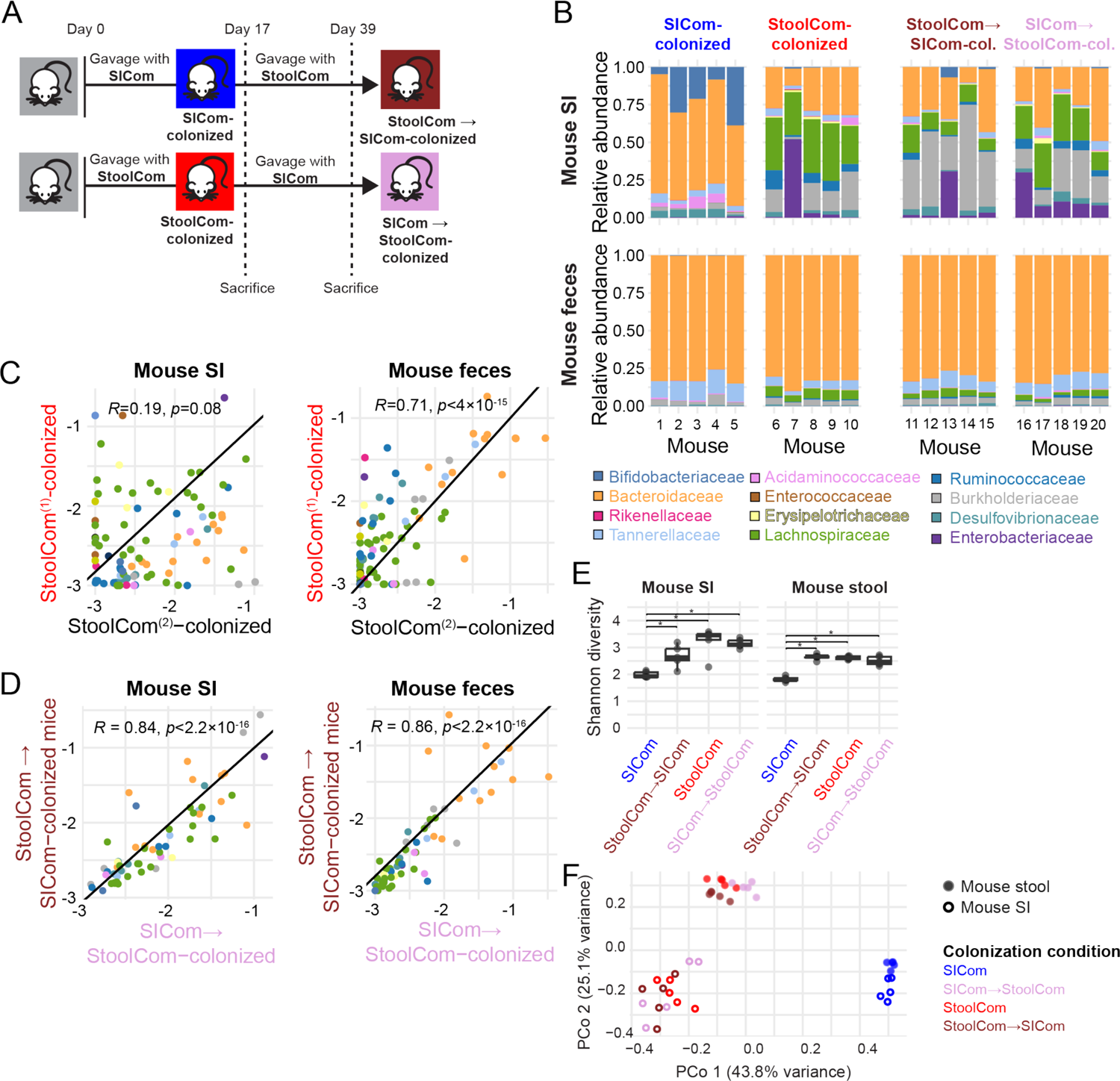
Reproducibility of mouse small intestine colonization is improved by the inclusion of human small intestinal microbes compared to human stool microbes alone. A) Two isolators each with two cages of five mice each were gavaged with SICom or StoolCom. After 17 days, one cage from each isolator was sacrificed and fecal and SI contents were collected, while the other cage was gavaged with the opposite community. After 39 days, all cross-colonized mice were sacrificed, and feces and SI contents collected. B) Family-level relative abundance of the SI (top) and fecal microbiotas (bottom) for SICom-, StoolCom^(2)^-, StoolCom→SICom-, and SICom→StoolCom-colonized mice (left to right). The SI microbiota of mice gavaged with both communities more closely resembled that of StoolCom^(2)^-colonized mice. Each bar represents an individual mouse. C) The fecal microbiota (left) exhibits a stronger correlation between StoolCom^(1)^- and StoolCom^(2)^-colonized mice compared to the SI microbiota (right). Each circle represents the mean log_10_(relative abundance) of an ASV, and color corresponds to family-level annotation. Shown are Pearson correlation coefficients. D) Fecal (left) and SI (right) microbiotas were highly correlated between cross-colonization conditions. Each circle represents the mean log_10_(relative abundance) of an ASV, and color corresponds to family-level annotation. Shown are Pearson correlation coefficients. E) Shannon diversity for mouse samples from the colonization conditions in (B-D). F) A Principal Coordinate Analysis showed that the composition of the mouse fecal and SI microbiota during cross-colonization conditions was more similar to that of StoolCom-colonized mice versus SICom-colonized mice.

For all colonization conditions, SI microbiota composition was highly distinct from that of feces (Fig. 3B, left). Like StoolCom-colonized mice, the SI microbiota of StoolCom^(2)^-colonized mice contained a high abundance of emergent ASVs (62.8%), while emergent ASVs accounted for only 6.1% of the SI microbiota of SICom-colonized mice. The large percentage of emergent ASVs in the SI of both cohorts of StoolCom-colonized mice led us to ask whether these emergent microbes are representative and colonize reproducibly. In fact, SI microbiota composition differed dramatically between the first cohort of StoolCom-colonized mice (henceforth, StoolCom^(1)^-colonized) and StoolCom^(2)^-colonized mice (Fig. 3C), suggesting that ASVs from StoolCom do not emerge reproducibly in the mouse SI, potentially due to their low abundance in the human stool-based *in vitro* community. Twenty-seven ASVs, accounting for 29% of the total abundance, and 3 families detected in the SI of StoolCom^(1)^-colonized mice were not detected in the SI of StoolCom^(2)^-colonized mice. Similarly, 17 ASVs, accounting for 28.5% of the abundance, and 4 families detected in the SI of StoolCom^(2)^-colonized mice were not detected in the SI of StoolCom^(1)^-colonized mice. By contrast, fecal microbiota composition was highly correlated between StoolCom^(1)^- and StoolCom^(2)^-colonized mice (Fig. 3C). Thus, colonization with a stool-derived *in vitro* community alone can lead to substantial stochasticity in SI colonization that is not apparent from analyses of feces, and may affect the reproducibility of other microbiota measurements and physiological outcomes.

Due to the distinct SI microbiota composition in SICom-colonized mice compared with StoolCom^(1,2)^- or DualCom-colonized mice, we wondered whether ordered colonization, enabling SI-specific microbes to engraft and gain priority over StoolCom microbes, would improve the reproducibility of SI microbiota colonization and limit the stochastic emergence of ASVs observed in StoolCom^(1,2)^-colonized mice. Seventeen days after gavage with either SICom or StoolCom, mice were gavaged with the other community (Fig. 3A, S3C,D). Fecal samples were collected periodically over an additional 22 days, after which mice were sacrificed and SI contents were collected. Both SI and fecal microbiota composition were similar between these two cross-colonization conditions (SICom→StoolCom-colonized and StoolCom→SICom-colonized; Fig. 3D). Unlike the comparison between StoolCom^(1)^- and StoolCom^(2)^-colonized mice, the same ASVs emerged from undetectable levels during cross-colonization with SICom and StoolCom. Furthermore, the SI microbiota composition of DualCom-colonized mice was significantly correlated with that of both SICom→StoolCom-colonized and StoolCom→SICom-colonized mice (*R*=0.66, *p*=8.6×10^-12^ and *R*=0.62, *p*=2.9×10^-10^, respectively; log_10_(ASV abundance), Pearson correlation). Thus, the introduction of SICom generally improves the reproducibility of SI microbiota composition across mouse experiments.

Despite the importance of SI microbes for the reproducibility of SI microbiota composition, several pieces of evidence argued against pervasive priority effects. Mouse feces contained the same families in the two cross-colonization conditions, as did the SI microbiota (Fig. 3B,D). The mouse fecal microbiota contained 66 ASVs shared across the two conditions, comprising >99.9% of the abundance in both StoolCom→SICom-colonized and SICom→StoolCom-colonized mice. Similarly, the SI microbiota contained 60 shared ASVs comprising 99.34% and 99.25% of the abundance for StoolCom→SICom-colonized and SICom→StoolCom-colonized mice, respectively. The composition of the mouse fecal and SI microbiota during cross-colonization was more similar to that of StoolCom-colonized mice versus SICom-colonized mice, suggesting that some species in StoolCom are dominant colonizers (Fig. 3E,F). In SICom→StoolCom-colonized mice, the family-level composition of the mouse SI microbiota was correlated with that of the human SI sample used to derive SICom (*R*=0.61, *p*=0.02; log_10_(family relative abundance), Pearson correlation) to a similar level as in StoolCom→SICom-colonized mice (*R*=0.56, *p*=0.037). Thus, the order of introduction has minimal effects on microbiota composition in either the SI or feces of mice.

### Assembly of a strain library representative of the SI microbiota

Isolates provide a powerful tool for illuminating how the physiology of a bacterial strain impacts community dynamics and structure. Furthermore, gnotobiotic mouse colonization using a synthetic community of isolates can circumvent reproducibility issues relating to emergent ASVs. To capitalize on these benefits, we utilized selective plating and FACS sorting^26^ to generate an extensive strain library, with the goal of encompassing a large fraction of the phylogenetic diversity observed in the human intestinal tract and stool. Using FACS, we isolated 109 strains directly from the device samples used to derive the *in vitro* communities described above and 406 strains from the *in vitro* communities.

Since the *in vitro* communities and device samples are undefined and contain strains at relative abundance below the limit of detection (<10^-3^), we reasoned that we could enrich for some of these strains by colonizing germ-free mice with SIcom or StoolCom. We sacrificed the mice after 17 days, collected SI contents and feces, and used selective plating to isolate 21 and 25 strains with distinct colony morphologies from the SI and feces, respectively. In total, these strategies resulted in 561 isolated strains.

We sequenced the V3 region of the 16S rRNA gene of these 561 isolates and found that they represent at least 64 species covering the major phyla of the human gut microbiota (Actinobacteria, Bacteroidetes, Firmicutes, and Proteobacteria) in 19 families according to BLAST annotation (Table S1). The main gut commensal from the Verrucomicrobiota phylum (*Akkermansia muciniphila*) was not isolated; it was not detected in any of the device samples or *in vitro* communities derived from this subject and hence may not be present.

Ultimately, the isolate library represents 58.3%-96.4% (96.4% for the sample used to generate SICom) of the total relative abundance in the intestinal samples from this subject and 69.7%-99.9% for the *in vitro* communities in Fig. 1F (99.8% for SICom and 76.7% for StoolCom). Interestingly, 3 of the 21 strains isolated from the SI of community-colonized mice and 18 of the 25 strains isolated from mouse feces did not overlap with the 515 strains isolated from device samples or *in vitro* communities, further highlighting that mouse colonization promotes the growth of certain species that are difficult to isolate directly from, or after *in vitro* culturing of, human gut samples.

We reasoned that these isolates were generally more likely to represent SI-specific versus stool-specific microbes. To enable construction of a synthetic community representative of stool, we isolated 31 strains using selective plating of a stool sample from the same subject that provided the device samples. Note that since all microbes in stool ultimately originated in the SI, these strains might be enriched in the SI despite isolation from stool. Based on 16S sequencing, these 31 strains represent at least 12 unique species from 3 phyla and 7 families. Of these stool-isolated strains, 9 species were not isolated in the SI-focused efforts described above, highlighting the distinct composition of the two regions.

To determine the extent to which our isolate library represents the human SI, we compared cultured strains to 16S sequencing of SI samples from 15 human subjects. Of the five taxa that were significantly enriched in the intestines across subjects^8^, three are represented by at least close relatives in the strain library (an *Escherichia* species, an *Enterococcus* species, and a *Bacteroides* (*Phocaeicola*) species closely related to *B. vulgatus*). For the remaining two taxa that were enriched in the human SI, the *Romboutsia* genus was undetectable in all samples and *Bilophila wadsworthia*, a fastidious species, was at <1.2% abundance in all *in vitro* communities, making isolation difficult (targeted isolation was not attempted from the original human intestine samples due to limited material).

Taken together, these isolates represent a phylogenetically diverse library from spatially distinct intestinal regions that is representative of most taxa in the human gut and *in vitro* communities (Fig. 4A).

**Figure 4:**
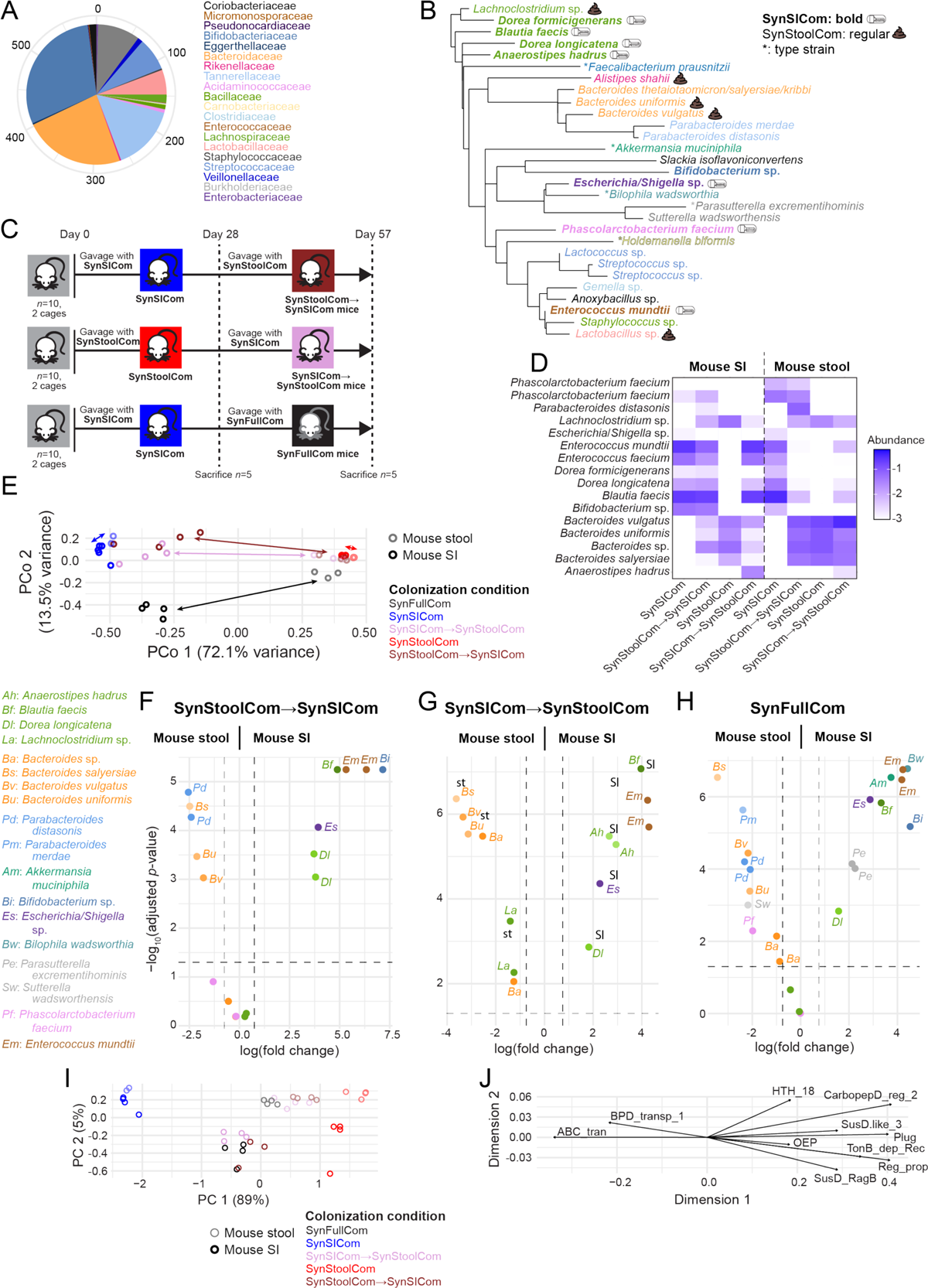
Synthetic communities of isolates recapitulate spatial niche preferences in mice. A) Family-level classification of a library of bacterial strains isolated from samples collected throughout the human intestinal tract and stool. B) A phylogenetic tree based on the ASVs of the strains used to construct synthetic communities SynSICom (bold), SynStoolCom, and SynFullCom that were gavaged into mice. The *Bacteroides thetaiotaomicron*, *salyersiae*, and *kribbi* strains were represented by the same ASV. Strains in SynSICom and SynStoolCom are denoted by a symbol after the name. Type strains included in SynFullCom that were used to supplement the library in (A) are denoted with an asterisk. C) Three isolators each with two cages of five germ-free mice each were gavaged with SynSICom or SynStoolCom. Twenty-eight days later, one cage of SynSICom- and SynStoolCom-colonized mice were sacrificed and feces and intestinal contents were collected. The remaining cages were gavaged with SynSICom, SynStoolCom, or SynFullCom as noted. After a total of 57 days, all mice were sacrificed and feces and intestinal contents were collected. D) Mouse microbiota composition was dependent on intestinal location and colonization condition. The mean log_10_(relative abundance) of each ASV is shown. E) Principal coordinate analysis based on Bray-Curtis distance between mouse microbiota compositions. All SI microbiota compositions were well separated from the corresponding fecal microbiota compositions except for SynSICom-colonized mice. Color denotes the colonization condition, and filled/empty circles denote stool/SI samples, respectively. F-H) ASVs with log_2_(fold-change)>0.75 and *p*<0.05 between mouse SI and stool are labelled with the species-level annotation for SynStoolCom→SynSICom (F), SynSICom→SynStoolCom (G), and SynFullCom-colonized mice (H). The R package limma-voom was used to calculate differential expression after size factors were estimated and normalized using DESeq2^50^. All *p*-values are Benjamini-Hochberg corrected. I) Principal component analysis (PCA) based on the relative abundance of Pfams in mouse intestinal microbiota metagenomes. Color denotes the colonization condition, and filled/empty circles denote stool/SI samples, respectively. J) The top 10 Pfams contributing to data stratification in the PCA in (I).

### Taxa maintain spatial preferences during synthetic community colonization

Synthetic communities of isolates are powerful tools that enable functional interrogation via whole genome sequencing of the isolates and follow-up *in vitro* experiments. Using our isolate library, we constructed three communities as potential models for the SI and fecal microbiota. Seven SI-derived isolates representing species enriched by >1.75-fold in the mouse SI compared to feces across four colonization conditions (StoolCom^(2)^, SICom, StoolCom→SICom, and SICom→StoolCom) were selected to represent the human SI microbiota (hereafter, SynSICom; Fig. 4B, Table S2, S3). To model the human stool microbiota, we selected seven species distinct from SynSICom that were (1) isolated from human stool or mouse feces or (2) present in mouse feces and not enriched in the mouse SI (hereafter, SynStoolCom; Fig. 4B, Table S3). Several other species that are prevalent across human gut microbiotas^27^ were not included in SynSICom or SynStoolCom, either because they were not isolated from the donor subject (due to low abundance or complete absence) or because they did not display a regional preference during mouse colonization with top-down *in vitro* communities. To account for the potential importance of these species in microbiota function, we also studied a phylogenetically diverse 31-strain community of SynSICom and SynStoolCom supplemented with *A. muciniphila*, which contributes to gut barrier function^28^, *B. wadsworthia*, an SI-enriched member associated with bile-acid deconjugation^8,29^, and several others (hereafter, SynFullCom; Fig. 4B, Table S3).

To interrogate whether these synthetic communities could reconstruct the site-specific colonization observed with the top-down *in vitro* communities SICom and StoolCom, we colonized separate cohorts of mice with SynSICom or SynStoolCom. On day 28, a subset of mice was sacrificed, and feces and SI contents were collected as described above. For the remaining mice, we introduced SynStoolCom into SynSICom-colonized mice, and vice versa, in separate isolators (Fig. 4C). On day 57, the cross-colonized mice were sacrificed, and feces and SI contents were collected and analyzed using 16S sequencing. For mice colonized with only SynStoolCom or SynSICom, the SI microbiota composition was very similar to that of the fecal microbiota (Fig. 4D,E), indicating spatial homogeneity of microbiota composition.

Cross-colonization with these synthetic communities revealed certain priority effects. Several observations supported the conclusion that when SynSICom colonization preceded SynStoolCom, aspects of regionality were generally better reconstituted. SynStoolCom member *Anaerostipes hadrus* was only detected in the mouse SI if it was introduced before SynSICom (Fig. 4D). Similarly, the *Bifidobacterium* species in SynSICom was only detected in the SI if it was introduced before SynStoolCom (Fig. 4D). SynStoolCom member *Parabacteroides merdae* was only abundant in the stool if it was introduced after SynSICom (Fig. 4C). Unlike cross-colonization involving SICom and StoolCom in which stool-derived microbes dominated microbiota composition throughout the intestinal tract (Fig. 3F), the SI microbiota composition of all cross-colonized mice was more similar to that of SynSICom-colonized mice compared to SynStoolCom-colonized mice (Fig. 4E). However, the fecal microbiota was similar to that of SynStoolCom-colonized mice across all cross-colonization conditions (Fig. 4E). Indeed, most members of SynSICom and SynStoolCom were significantly enriched in particular intestinal regions (Fig. 4F,G), suggesting that gut commensals have a geographical preference along the intestinal tract driven by local ecological factors. For instance, an *Escherichia/Shigella* species, *Dorea longicatena,* and a *Bifidobacterium* species isolated from the upper human intestinal tract were enriched in the mouse SI, and *Bacteroides salyersiae, B. thetaiotaomicron, Bacteroides kribbi,* and *Parabacteroides distasonis* isolated from mouse or human stool were consistently enriched in feces. Interestingly, although *Anaerostipes hadrus, Blautia faecis,* and *Enterococcus mundtii* were isolated from mouse or human stool, they were consistently enriched in the SI during colonization with *in vitro* or synthetic communities (Fig. 4F,G). By contrast, *Bacteroides uniformis* and *B. vulgatus* isolated from a human upper intestinal sample were consistently enriched in mouse feces (Fig. 4F,G). Thus, regardless of region of isolation, taxa typically exhibited consistent regional specificity in mice.

In two cases, a species isolated from human stool colonized the mouse SI and feces promiscuously. A *Lachnoclostridium* species was enriched in the mouse SI during certain colonization conditions with top-down *in vitro* communities but was assigned to SynStoolCom based on its isolation origin as well as the motivation to include a Firmicute with the potential for stool localization. This species was significantly enriched in stool when SynSICom was introduced to SynStoolCom-colonized mice, but not vice versa (Fig. 4F,G). Meanwhile a *Bacteroides* species closely related to *B. thetaiotaomicron* was significantly enriched in mouse feces when SynSICom was introduced to SynStoolCom-colonized mice but not vice versa (Fig. 4F,G). Thus, the region from which a species is isolated, priority effects, and community context can collectively determine localization preference in the intestinal tract.

Seventeen days after SynSICom colonization, we introduced SynFullCom to a subset of mice (Fig. 4C). These mice were sacrificed after an additional 22 days, and stool and SI contents were collected. Most SynFullCom members not in SynSICom or SynStoolCom displayed strong regional preferences, including localization of *A. muciniphila* and *B. wadsworthia* to the SI (Fig. 4H). Morever, almost all SynSICom and SynStoolCom taxa maintained their expected regional preference (Fig. 4H); the lone exception was *Phascolarctobacterium faecium*, which was at very low abundance (0.2%-1.8% across all intestinal regions). Thus, regional specificity is largely robust to challenge.

### Region-specific microbiotas have distinct functional capacities

Given the variation in microbiota composition across the mouse intestinal tract and the consistent regional specificity of certain species, we sought to quantify functional differences between SI- and stool-localized species. We sequenced the genomes of all isolates in the synthetic communities, and performed metagenomic sequencing of all mouse samples (intestinal and feces) involving colonization with synthetic communities. From these data, we assembled genomes, identified putative genes, and annotated their function using protein domain information from the Pfam database^30^ (Methods).

We calculated the functional dissimilarity between sample pairs based on the vector of individual Pfam counts normalized to the total number of Pfam counts in each sample. Similar to taxonomic abundances, SynSICom- and SynStoolCom-colonized mice displayed the most dissimilar Pfam profiles in both the SI and stool. However, unlike taxonomic abundances, for which cross-colonized mice exhibited SI and fecal microbiota composition similar to that of SynSICom- and SynStoolCom-colonized mice, respectively (Fig. 4E), the Pfam profile of cross-colonized mice was intermediate between SynSICom- and SynStoolCom-colonized mice in both the SI and feces (Fig. 4I). These findings indicate that introduction of SI microbes provides functional capabilities distinct from those of even closely related stool microbes and that 16S sequencing alone may be insufficient for determining the complete physiological impact of SI microbes.

We next sought to identify functional categories driving regional microbiota differences. Of the 10 Pfam domains that were the most important drivers of functional differences (Fig. 4J, Methods), eight are associated with substrate import and export, including channels, secreted proteins that bind substrates, and plugs that block channels. The other two domains are regulators, one of unknown function and one involved with L-arabinose utilization. The binding protein-dependent transport system inner membrane component (BPD_transp_1) and ABC transporter domains were responsible for separating SynSICom-colonized mice from other colonization states. The domains driving functional differences between the SI and fecal microbiota in cross-colonized states involve trade-offs between regulation of L-arabinose utilization (HTH_18) in the feces and RagB (similar to SusD, an OM-tethered protein that binds to glycans) in the SI (Fig. 4J). Thus, method of substrate transport and functional potential for substrate breakdown and import drive functional differences along the intestinal tract and between colonization states.

In SD-fed SynFullCom-colonized mice, 5,684 and 7,160 unique Pfams were identified in the feces and SI, respectively. Functional overlap between the two regions was very high, with only 80 Pfams detected solely in stool and 1,556 detected solely in the SI. These 1,556 SI-specific Pfams accounted for 0.09-1.7% of the relative abundance of Pfams in the SI. The 40 of these 1,556 Pfams detected in the SI of all mice include functions associated with phage and steroid (e.g., bile acid) processing, consistent with our previous observations that phage induction and bile acids are more abundant in the human SI^8^, suggesting differential selective pressures along the intestinal tract.

### A fiber-deficient diet causes a strain-dependent microbiota shift associated with isolation origin

To assess the role of MAC processing on microbiota composition in mice colonized with our synthetic communities, we switched a cohort of SynFullCom-colonized mice from a SD to a MD 15 days after colonization (Fig. 5A). Fourteen days later, the mice were sacrificed, and feces and intestinal contents were collected and analyzed. The diet switch caused taxonomic changes throughout the mouse intestinal tract (Fig. 5B,C). Altogether, a MD resulted in dramatic changes to fecal and SI microbiota composition, with colonization condition also having a small effect (Fig. 5B,C).

**Figure 5:**
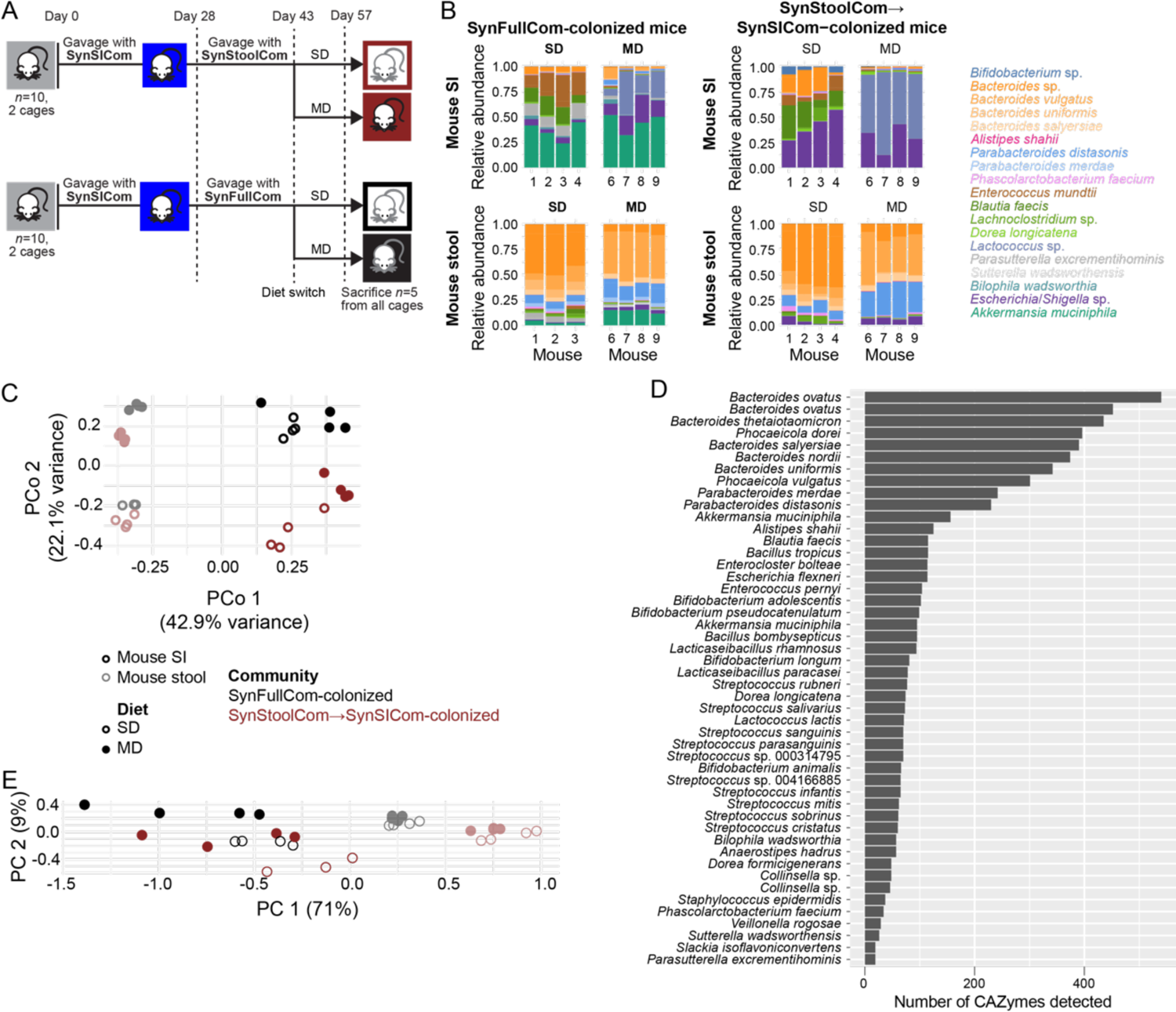
Synthetic communities exhibit diet-dependent composition and function across the intestines of mice. A) In two isolators each with two cages of five mice each, all mice were gavaged with SynSICom. After 28 days, one isolator was gavaged with SynStoolCom and the other was gavaged with SynFullCom. After 15 days, mice were either maintained on a standard diet (SD) or switched to a MAC-deficient diet (MD). After 57 days total, mice were sacrificed and intestinal and fecal contents were collected. B) ASV-level relative abundance was highly distinct between the mouse SI (top) and fecal (bottom) microbiotas for SynFullCom-colonized-(left group) and SynStool→SynSICom-colonized (right group) mice. Within each colonization condition, the diet switch from a SD to a MD alters both the SI and fecal microbiotas. Each bar represents an individual mouse. C) Principal coordinate analysis based on Bray-Curtis distance between mouse microbiotas based on the relative abundance of ASVs. Each light/dark circle represents a mouse fecal/SI microbiota, respectively. The color denotes the colonization condition, and empty/filled circles indicate a SD/MD diet, respectively. D) The number of CAZymes detected from whole genome sequencing varied extensively across a subset of the collection of bacterial isolates acquired from throughout the intestinal tract shown in Fig. 4A (Methods). E) Principal component analysis (PCA) based on the relative abundance of Pfams between mouse intestinal microbiotas separates SI microbiotas by diet and colonization condition. Each light/dark circle represents a mouse fecal/SI microbiota, respectively. The color denotes the colonization condition, and empty/filled circles indicate a SD/MD diet, respectively.

In feces, the relative abundance of MAC-degrading *B. thetaiotaomicron* decreased and that of mucus-degrading *A. muciniphila* increased, as expected based on nutritional preferences^31^ (Fig. 5B). The diet switch also resulted in an increase in the relative abundance of *B. vulgatus* and *P. distasonis* (Fig. 5B), consistent with their lower proportion of carbohydrate active enzymes (CAZymes) compared to *B. thetaiotaomicron* (Fig. 5D). These changes are consistent with our findings for StoolCom^(1)^- and DualCom-colonized mice on a MD diet (Fig. 2C). However, the diet switch increased the abundance of *Alistipes shahii* in the fecal microbiota of StoolCom^(1)^-colonized mice (Fig. 2C) but not SynFullCom-colonized mice (Fig. 5B), potentially because the *A. shahii* in SynFullCom was isolated from the human intestinal tract rather than stool. This observation suggests the potential for strain-dependent responses to diet changes that may be driven by regional preference.

In the SI, the relative abundance of *A. muciniphila* did not increase in MD-fed relative to SD-fed mice, likely due to its already high relative abundance in the SI of SD-fed mice. Instead, the relative abundance of *Lactococcus lactis* increased significantly and that of MAC-degrading *Blautia faecis* decreased (Fig. 5B,D). In SynFullCom-colonized mice, the relative abundance of the *Enterococcus* species, which was isolated from human stool, was dramatically lower in the SI of MD-fed compared to SD-fed mice (Fig. 5B). This behavior was unexpected based on mice colonized with top-down *in vitro* communities. In StoolCom^(1)^-colonized mice, the relative abundances of two *Enterococcus* ASVs closely related to the *Enterococcus* isolate in our synthetic communities were unaffected by a diet shift (Fig. 2I). By contrast, in the SI of DualCom-colonized mice, the relative abundance of those *Enterococcus* ASVs (as well as a third *Enterococcus* ASV) all increased significantly after the diet switch, suggesting that the same ASV may represent multiple strains with distinct responses to diet.

We next assessed how the functional potential of the microbiota is altered by a MD using Pfam annotations as above. The Pfam profile in the SI showed more variation with respect to diet in both SynFullCom- and SynStoolCom→SynSICom-colonized mice than in the feces (Fig. 5E). In contrast to compositional data (Fig. 5C), fecal microbiota Pfam abundance was more strongly affected by colonization condition than by diet (Fig. 5E). Thus, the taxonomic changes due to a diet shift are not necessarily associated with altered functional potential. However, SI microbiota Pfam abundance showed a strong dependence on diet as well as composition (Fig. 5E). Taken together, functional alterations to the microbiota due to a MD can be more apparent in the SI than in feces.

To determine if these different behaviors reflect strain-specific metabolic differences, we isolated two and three *Enterococcus* strains from the SI of MD-fed StoolCom^(1)^- and DualCom-colonized mice, respectively, and sequenced their genomes along with that of the *Enterococcus* strain from our synthetic communities. Four of the strains (the two from StoolCom and two of the three from DualCom) were identified as *E. faecium*, and were essentially identical (with average nucleotide identity (ANI) of 100). The synthetic community strain, which was isolated from the human stool sample, was identified as *E. mundtii*, and the third DualCom strain was identified as *E. avium*; for both strains, the core genome had an ANI relative to *E. faecium* of ∼75-80. The *E. avium* strain was the closest match to the *Enterococcus* ASV that bloomed after the switch to a MD and reached the highest abundance in the SI of DualCom-colonized mice. To investigate potential differences in metabolic capacities that might underlie their responses to the MD diet switch, we analyzed the CAZymes encoded in their genomes and the genomes of a clinical isolate of *E. avium* and the type strain *E. mundtii* DSM4838 (Methods). As expected, all *E. faecium* strains had identical CAZyme profiles, which were highly similar to that of a type strain DSM20477 (Fig. 6A). The *E. avium* strain encoded 15 CAZymes families that were almost entirely absent from the *E. faecium* strains and *E. mundtii*.

**Figure 6:**
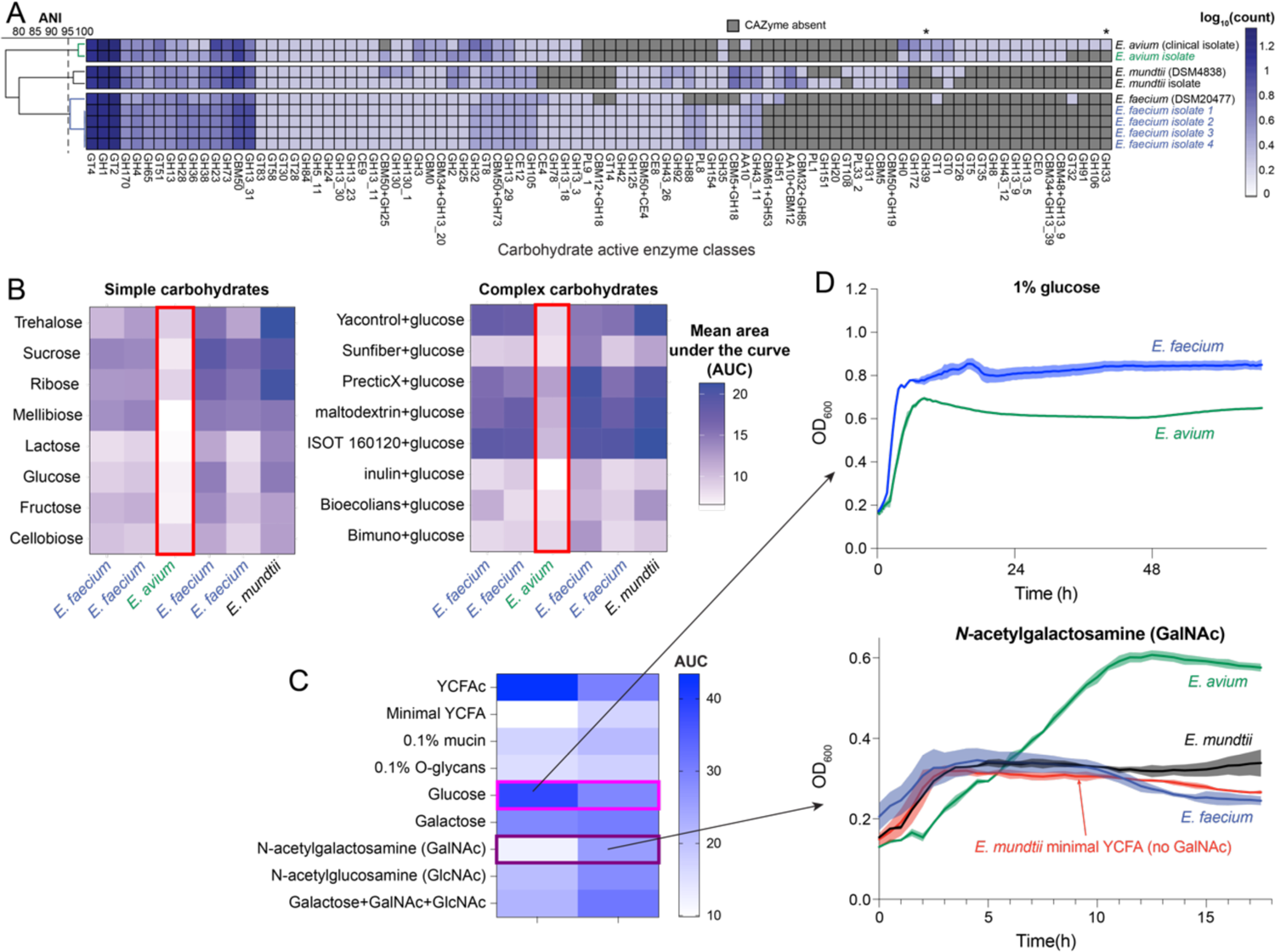
Blooming of *Enterococcus avium* in the SI of mice fed a MD is associated with the ability to utilize mucin components. A) Carbohydrate active enzymes (CAZymes) detected from whole genome sequencing of our *Enterococcus* isolates and representative clinical isolates or type strains. Shown on the left is a phylogenetic tree based on MASH average nucleotide identity (ANI). Asterisks at the top denote the presence of GH39 CAZymes uniquely in the *E. avium* strains and the presence of GH33 sialidases involved in mucin utilization uniquely in the *E. avium* clinical isolate. B) *Enterococcus* growth on carbohydrates varied across isolates. The *E. avium* strain isolated from DualCom-colonized mice was unable to grow well on any of the simple or complex carbohydrates tested (red boxes). This *E. avium* strain was the closest match to the ASV that bloomed after the switch to a MD in the SI of DualCom-colonized mice (Fig. 2G). Area under the curve (AUC) was quantified as an average across four replicate growth curves after 48 h of growth in minimal medium + 0.25% of the indicated carbohydrate. C,D) Compared with a representative *E. faecium* isolate, the *E. avium* isolate grew worse in glucose (maroon box) but was uniquely able to utilize the mucin component N-acetylgalactosamine (GalNAc, purple) among the *Enterococcus* strains.

In contrast to the rest of the *Enterococcus* strains (Fig. 6B), *E. avium* grew little if at all on a wide range of complex carbohydrates and grew slowly in disaccharides such as sucrose and trehalose (Fig. 6B), consistent with its lower fitness in SD-fed mice. Based on the expansion *of E. avium* in a MD diet, during which enhanced mucin utilization is expected^16,20^, we hypothesized that *E. avium* would uniquely have expanded capacity for utilization of host-derived nutrients such as mucins. Although some *Enterococcus* strains have been reported to degrade host mucins^32^, we found that all *Enterococcus* strains were unable to consume intact mucin or purified O-glycans; this finding was consistent with the lack of glycoside hydrolase (GH) 33 sialidases in the genome of all of our *Enterococcus* strains, despite the presence of GH33 enzymes in a clinical isolate of *E. avium* (Fig. 6A). *E. faecium* exhibited a higher maximum growth rate and yield than *E. avium* in minimal YCFA (mYCFA) with supplemented glucose (Fig. 6C). However, *E. avium* was able to utilize GalNAc, a major component of glycans found in mucus, and grew to a similar yield as *E. faecium* in glucose, while *E. faecium* and *E. mundtii* were unable to grow beyond the background growth observed in the mYCFA base medium (Fig. 6D). *E. avium* also exhibited more growth than *E. faecium* in GlcNAc, another monosaccharide component of mucus glycans (Fig. 6C, S4). Thus, the variable responses of these strains in the SI during a MD are consistent with differences in mucin-utilization capabilities. Furthermore, these findings indicate that gavage with SI-derived communities introduced closely related but metabolically distinct species, including those from the *Enterococcus* genus, highlighting the importance of inoculating with SI microbes when studying intestinal biology.

## Discussion

The variation in physiology and function along the human intestinal tract provides a wide range of niches conducive to microbiota colonization. As a result, individual members of the microbiota have been proposed to have closely matched physiology and genetic potential to the region they occupy. Stool samples, while easily accessible, contain a mixture of microbes from throughout the intestinal tract with limited information about their regional preferences. In this study, we applied a multifaceted approach to demonstrate that colonization with cultured human SI-derived microbial communities improves humanized gnotobiotic mouse models for the SI microbiota and provides novel information about the ecology of the SI microbiota. By capturing spatially distinct microbial communities using a noninvasive sampling device, we established stable, reproducible, and representative communities of SI microbiota^8,13,33^ (Fig. 1). We found that colonization with a combination of SI-and stool-derived communities led to distinct, more reproducible SI microbiota compositions (Fig. 3C,D) and altered the response to dietary changes relative to a stool-derived community alone (Fig. 2C,F-I). Thus, future experiments with gnotobiotic or humanized mice should consider the importance of SI microbes, for which our isolate library, top-down *in vitro* communities, and bottom-up synthetic communities can serve as useful tools.

Cultivation of intestinal contents proved critical for probing SI biology by enabling reproducible inoculation of microbes representative of the intestinal tract at the volumes necessary for gavaging a cohort of mice. Since the SI microbiota varies along the intestinal tract and across human subjects^8^, generating communities and isolates from a variety of spatially distinct regions and from multiple humans (both healthy and diseased patients) should enable further interrogation of SI biology, especially regarding the role of prevalent, SI-enriched genera such as *Romboutsia* and *Bilophila* that happened to be absent or in low abundance in the subject used in this study.

By assembling a strain library representative of the SI microbiota, we constructed phylogenetically diverse synthetic communities and identified taxa that maintain a spatial preference in mice (Fig. 4F-H) and whose response to a diet switch can be dependent on isolate source (Fig. 2C,G, 5B). We further profiled functional differences in the microbiota along the intestinal tract and uncovered region-specific functional changes in response to diet (Fig. 5F). While the colon is primarily exposed to non-host-absorbed components of diet, the SI is exposed to varying degrees of dietary components, from the duodenum, which sees nearly all dietary intake, to the ileum, which has less exposure to simple carbohydrates and other substances absorbed more proximally^34,35^. Our finding that fiber-utilizing bacteria are depleted within the SI during a fiber-free diet (Fig. 2C,G, 5B) suggests that MACs can be fermented in the SI and support microbiota structure at this site. Furthermore, the larger decrease in SI versus fecal microbiota diversity in response to the diet switch (Fig. 2F) has important implications for understanding community recovery in response to diet alterations.

Germ-free mice humanized with human stool have proven powerful for facilitating mechanistic understanding of the human gut microbiota, but have many caveats and limitations, including incomplete colonization of SI-resident bacteria^36,37^. Our study demonstrates that human gut commensals largely exhibit similar taxonomic enrichment between the mouse SI and feces as between the human SI and stool^8^, supporting the relevance of mice as a model for the human intestines. However, mice did not exhibit the degree of variation between regions of the SI (duodenum, jejunum, and ileum) observed in humans^7^. We speculate that this difference may be due to the dietary homogeneity and continuous eating patterns of mice, which could be addressed by future studies of wild mice. Coprophagy may also partially homogenize the small intestine by providing additional colonization opportunities for SI microbes.

Although we demonstrated that gavage with SI microbes can stabilize colonization of the SI (Fig. 3C,D), the degree to which small intestinal niches are filled remains undetermined. We observed increased priority effects involving species at <1% abundance using synthetic communities with reduced diversity relative to top-down *in vitro* communities (Fig. 5B), which has implications for microbiota recovery after a disturbance that dramatically reduces diversity such as antibiotics or infections. In the future, strategies based on ecological back-filling^15^ could be used to determine if additional exposure to SI microbes will further increase richness in the SI and if strain replacement occurs, which are important considerations for therapies targeting the SI microbiota. Given that SI microbiota dysbiosis is emerging as a hallmark of metabolic disorders as well as functional and inflammatory gastrointestinal disease in humans^38–40^, our study provides a foundation for developing improved animal models that recapitulate such dysbiosis, to test whether SI-derived microbiota from healthy individuals can colonize and to reverse microbiota alterations in various disorders by directly targeting small intestinal ecology^6,41,42^.

Our identification of site-specific bacterial colonization and regionally distinct responses to diet highlights the need to further understand the spatial biogeography and functional heterogeneity along the length of the gastrointestinal tract, which should complement both animal models and *in vitro* models of the intestinal ecosystem^43^. The specific expansion of *E. avium* in the small intestine of mice fed a fiber-deficient diet is consistent with both the inability of this isolate to process complex polysaccharides (Fig. 6B) and its unique capacity to utilize mucus components (Fig. 6C,D). The inability of all our *Enterococcus* isolates to utilize mucins and O-glycans, presumably due to their lack of certain GH activities required to de-cap or for processive monosaccharide removal such as the GH33 sialidases (Fig. 6A)^44^, suggests that other species facilitate *E. avium* expansion. Similarly, *E. coli* is well known to utilize monosaccharides found in mucus even though it does not possess the GHs to liberate these sugars^45^. While we identified several bacterial species with preferential SI colonization and distinct functional properties, specific mechanisms that mediate regional specificity within the intestine remain unclear. These large knowledge gaps motivate future large-scale efforts to isolate and characterize bacteria collected from the human SI. Collection of microbiota samples from the SI, via capsule devices or endoscopically, is laborious and provides only limited biomass within samples. Our mouse experiments revealed that SI-colonizing bacteria within stool samples can be isolated by leveraging gnotobiotic mice as a culturing platform. Moreover, disease-associated bacteria are sometimes found at low abundance in stool but enriched in SI samples, suggesting that these species are primarily resident in the SI and impact host physiology there^8,29^. These points demonstrate the importance of developing gnotobiotic mouse models to understand region-specific microbe-host interactions.

Identification of species in stool with preference for the SI will provide greater mechanistic insight into the extensive microbiome studies involving stool. Future microbiota-targeted therapeutic approaches using live biotherapeutic products will be enhanced by inclusion of SI-derived strains targeted for SI colonization. Moreover, fecal microbiota transplant (FMT) delivery directly to the SI could lead to miscolonization of stool-resident bacteria into small intestinal niches, particularly in patients preconditioned with niche-clearing antibiotics and receiving treatment via duodenal instillation. The regional variation in microbiota composition after FMTs can be tested in humans using samples obtained from our capsule devices and should be considered as a factor that can either enhance or reduce FMT therapeutic efficacy and as a source of the variability in clinical trials due to the content of SI microbes and their exposure to different gut regions based on delivery method (e.g., colonic delivery versus duodenal instillation).

Overall, these findings represent a significant advance in our ability to study and understand host-microbe dynamics in the SI. As the SI is the site of dietary absorption, SI-resident microbes have a large impact on nutrient uptake and may provide signals that promote barrier function and metabolic homeostasis in health and may perpetuate inflammation and metabolic disease when altered^11^. Our study will enable development of humanized gnotobiotic mouse models, potentially even personalized mouse avatars colonized with patient-derived microbiota, that better recapitulate human disease states and enable testing of SI-targeted microbial therapeutics.

## Methods

### Mice

All mouse experiments were conducted in accordance with the Administrative Panel on Laboratory Animal Care, Stanford University’s IACUC. Germ-free, Swiss-Webster mice (male and female) were initially colonized with either 200 µL of a small intestine-derived or stool-derived community grown for 48 h, or a collection of isolates grown separately for 72 h and then mixed at equal OD on the day of gavage. Mice inoculated with different microbial communities were kept in separate isolators. Mice with the same colonization condition but fed different diets were kept in separate cages within the same isolator. Fecal pellets were sampled periodically and frozen immediately.

Mice were euthanized via cervical dislocation. Fecal pellets were collected immediately before death. The small intestine was dissected and folded into three equally sized sections that were collected in order (proximal, middle, and distal). Three milliliters of sterile PBS were used to flush the intestinal contents out of each section into a tube. Cecal contents were dissected and collected into a tube. A ∼350 µL aliquot of each small intestine or cecal sample was added individually to 150 µL of sterile 50% glycerol for future isolation experiments before freezing.

### In vitro community passaging

To generate SI-derived *in vitro* communities, a single subject swallowed capsule devices that sampled from various regions of the intestinal tract^8^. Samples were acquired after the necessary approval from the Institutional Review Board (IRB), and informed consent was duly obtained from the subject prior to sample collection. Each device was quickly stored in nitrogen after collection to establish an anaerobic environment and preserve the viability of the contents. An aliquot of stool was also sampled and preserved in a similar manner. Contents were then transferred to an anerobic chamber. Capsule samples were extracted using sterile needles and ∼100 mg of stool were resuspended in 500 µL of sterile, pre-reduced PBS and vortexed vigorously. Approximately 50 µL of contents were set aside for sequencing, and 10 µL were added to glycerol for storage at −80 °C.

One microliter of each sample was used to inoculate 200 µL of pre-reduced medium in a 96-well microplate. Each culture was passaged repeatedly with 1:100 dilutions every 48 h at 37 °C in an anaerobic chamber. Each community was passaged in triplicate in Brain Heart Infusion (BHI) at various pH values and in various concentrations of the pH-appropriate buffer. After 14 days (7 passages), communities were mixed with glycerol (final concentration 15%) for storage at −80 °C.

### Bacterial strain isolation

Strains were isolated using either plating from *in vitro* communities or capsule devices or sorting from *in vitro* communities (Table S1). For plating, communities or capsule device contents were plated in ten-fold dilutions onto Brain Heart Infusion (BHI)+10% blood and incubated for 72 h at 37 °C in an anaerobic chamber. Single colonies were picked using sterile toothpicks and re-streaked onto BHI+10% blood plates. This process was repeated an additional two times to ensure that the colony was axenic. Single colonies were then picked into a 2-mL deep-well plate containing 500 µL of BHI supplemented with menadione (vitamin K), cysteine, and hemin (BHIS) or Reinforced Clostridial Medium supplemented with menadione, cysteine, and hemin (RCMS).

Some isolates were also acquired by sorting *in vitro* communities into BHIS using a cell sorter as previously described^26^. After 72 h of growth in an anaerobic chamber at 37 °C, isolates were mixed with glycerol (final concentration of 12%) and stored at −80 °C.

To determine the species identity of isolates, we conducted Sanger sequencing. Cultures were spun down and resuspended with PCR-grade water in a 1:1 ratio. The primers 5’AGAGTTTGATCCTGGCTCAG and 5’GACGGGCGGTGWGTRCA were used as previously described^46^. Taxonomic assignment was performed using NCBI BLAST.

The library of 561 isolates in this paper was condensed to one 96 well plate, referred to as the Small Intestine Isolate LibrarY (SIILY), as follows. All target isolates were struck onto BHI+10% blood plates. After 72 h, single colonies were picked into a 96 deep-well plate containing RCMS. Certain known fastidious strains were additionally supplemented with nutrients: *Bilophila wadsworthia* with 4 mM taurine, *Akkermansia* strains with 0.01% mucin, *Veillonella* strains with 1.5% lactate, and *Phascolarctobacterium faecium* with 0.2% succinate. After 72 h, cultures were diluted 1:10 and final OD_600_ was measured using an Epoch 2 plate reader (Biotek). Isolates were mixed with glycerol (final concentration of 12%) and stored at −80 °C.

### High-throughput measurements of growth dynamics

Optical density at 600 nm (OD_600_) measurements at ∼3 min intervals were performed with an Epoch 2 Biotek plate reader in an anaerobic chamber at 37 °C with continuous shaking. Growth curves were measured in a 96-well plate for 48-72 h for isolates and 48 h for *in vitro* communities. To measure pH, BCECF dye (ThermoFisher, B1150) was used.

### 16S rRNA gene sequencing

DNA was extracted from whole fecal pellets or 50 µL of *in vitro* cultures with a DNeasy PowerSoil HTP 96 kit (Qiagen 12955-4) or a DNeasy UltraClean Microbial Kit (Qiagen 10196-4). 16S rRNA gene amplicons were generated using Earth Microbiome Project-recommended 515F.806R primer pairs using the 5PRIME HotMasterMix (Quantabio 2200410) as previously described^46^. This product was run on a 1% TAE gel, and a band at ∼300 bp was gel extracted. Raw data were demultiplexed using QIIME2^47^. Filtering and trimming were conducted using DADA2^48^ as previously described^33^. Resulting ASV sequences were assigned taxonomy using the SILVA database^49^ and assignTaxonomy() and addSpecies() from DADA2^48^.

### Differential enrichment

To determine ASVs whose abundance was significantly different between the SI versus feces, the R package limma-voom was used to calculate differential expression after size factors were estimated and normalized using DESeq2^50^.

### Metagenomic and whole genome sequencing

Extracted DNA was arrayed into a 384-well plate and concentrations were quantified using the PicoGreen dsDNA quantitation kit (ThermoFisher) for normalization. DNA wells were individually tagmented and barcoded using the Nextera XT kit (FC-131-1096). One microliter from each well was mixed, and the pooled library was purified twice using Ampure XP beads to select the appropriately sized bands. Finally, library concentration was quantified using a Fragment Analyzer. Isolate sequencing was performed on a HiSeq 4000 with read lengths of 2×150 bp. Metagenomic sequencing was performed on a NovaSeq S4 with read lengths of 2×150 bp.

Skewer v. 0.2.2^51^ was used to remove Illumina adapters. Reads were assembled with SPAdes v. 3.15.5^52^. checkM v. 1.1.3^53^ and quast v. 5.0.2^54^ were used to assess quality. All genomes were de-replicated at 99% ANI (strain level) with dRep v. 3.0.0^55^ to generate the 48 unique species whose genomes were analyzed for their CAZyme abundances (Fig. 5D). GTDB-Tk was used to assign taxonomy^56^. Default parameters were used for all computational tools.

To calculate average nucleotide identity (ANI) between *Enterococcus* genomes, dRep v. 3.4.5^55^ was used with the following parameters: compare, -ms 100000, - pa 0.95. Putative genes were called on assembled contigs for each sample or on assembled MAGs using Prodigal^57^. CAZyme genes were identified using run_dbcan.py v. 3.0.5 or v. 4.1.4^58^ using default parameters (searching via HMMER, eCAMI, and DIAMOND). Genes identified in at least 2 of 3 programs were utilized in subsequent analyses. Prodigal^57^ was also used to determine putative proteins, and HMMER was used to annotated Pfams. Annotation results from the DIAMOND search against the CAZy database were summarized for subsequent analysis.

### Enterococcus growth dynamics

Isolated *Enterococcus* strains were struck onto BHI agar plates and incubated for 24 h. A colony from each plate was inoculated in YCFAc (Yeast Casitone Fatty Acids Agar with carbohydrates) broth for 24 h. One hundred eighty microliters of each mYCFA solution with different carbon source additives was mixed with 20 µL of an *Enterococcus* strain in each well of a 96-well plate. Optical density at 600 nm (OD_600_) measurements at 37 °C with continuous shaking were performed at 30 min intervals with an Epoch 2 Biotek plate reader in an anaerobic chamber.

## Data availability statement

All sequencing data is available is available on NCBI Sequencing Read Archive under Project PRJNA948701 (16S datasets: SRA PRJNA948701). A github repository containing R v. 4.0^59^ scripts for the generation of all figures and analyses is available at https://github.com/rculver123/modeling-small-intestine-microbiota.git.

## Acknowledgements

The authors thank members of the Huang lab for helpful discussions, and acknowledge support from a National Defense Science and Engineering Graduate Fellowship (to R.N.C.), the Colleen and Robert D. Hass fund (S.P.S.), NIH grants RM1 GM135102 (to K.C.H. and J.S.), R01 AI147023 (to K.C.H.) T32DK007056 (to S.P.S.), and K08DK134856 (to S.P.S.), and NSF Grants 2126329 (to D.S.), and EF-2125383 and IOS-2032985 (to K.C.H.). K.C.H. and J.S. are Chan Zuckerberg Biohub Investigators.

## Supplementary Figures

**Figure S1:**
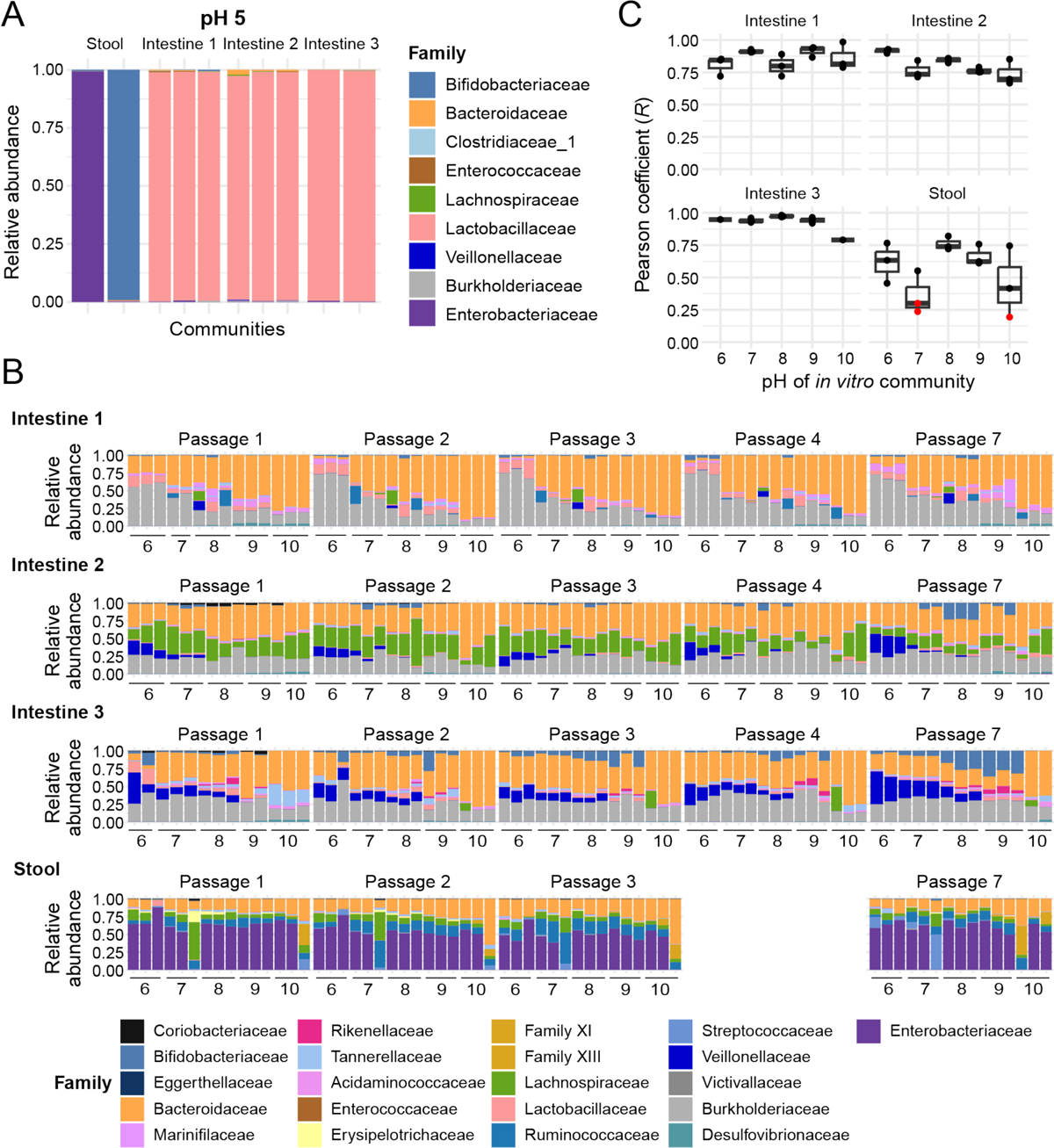
*In vitro* community growth was sensitive to low pH, equilibrated quickly, and was highly reproducible. A) *In vitro* communities derived from stool grown at pH 5 were dominated by the Enterobacteriaceae or Bifidobacteriaceae family, while communities derived from intestinal samples were dominated by a single *Lactobacillus* species. B) *In vitro* community composition equilibrated quickly. C) Technical replicate *in vitro* communities generally reached highly similar compositions across starting pH values, particularly for intestinal samples.

**Figure S2:**
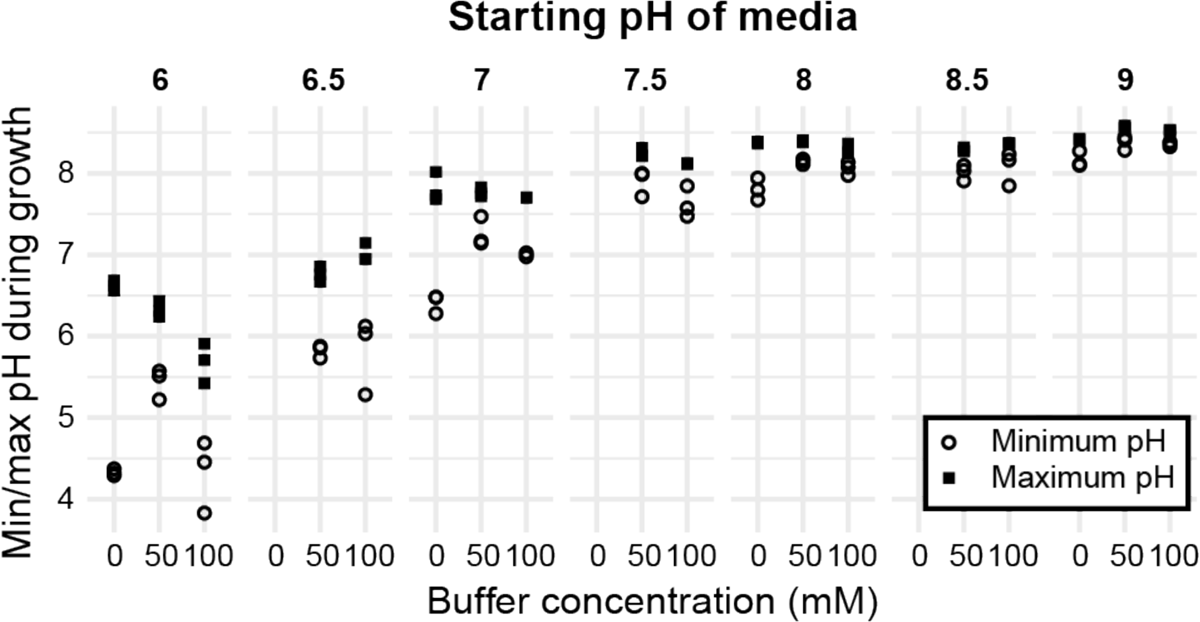
Buffer addition restricts modification of environmental pH. Shown are the minimum (circles) and maximum (squares) pH values for *in vitro* communities grown in BHI with the starting pH value shown above each plot, with 0, 50, or 100 mM of the buffer appropriate for the starting pH.

**Figure S3:**
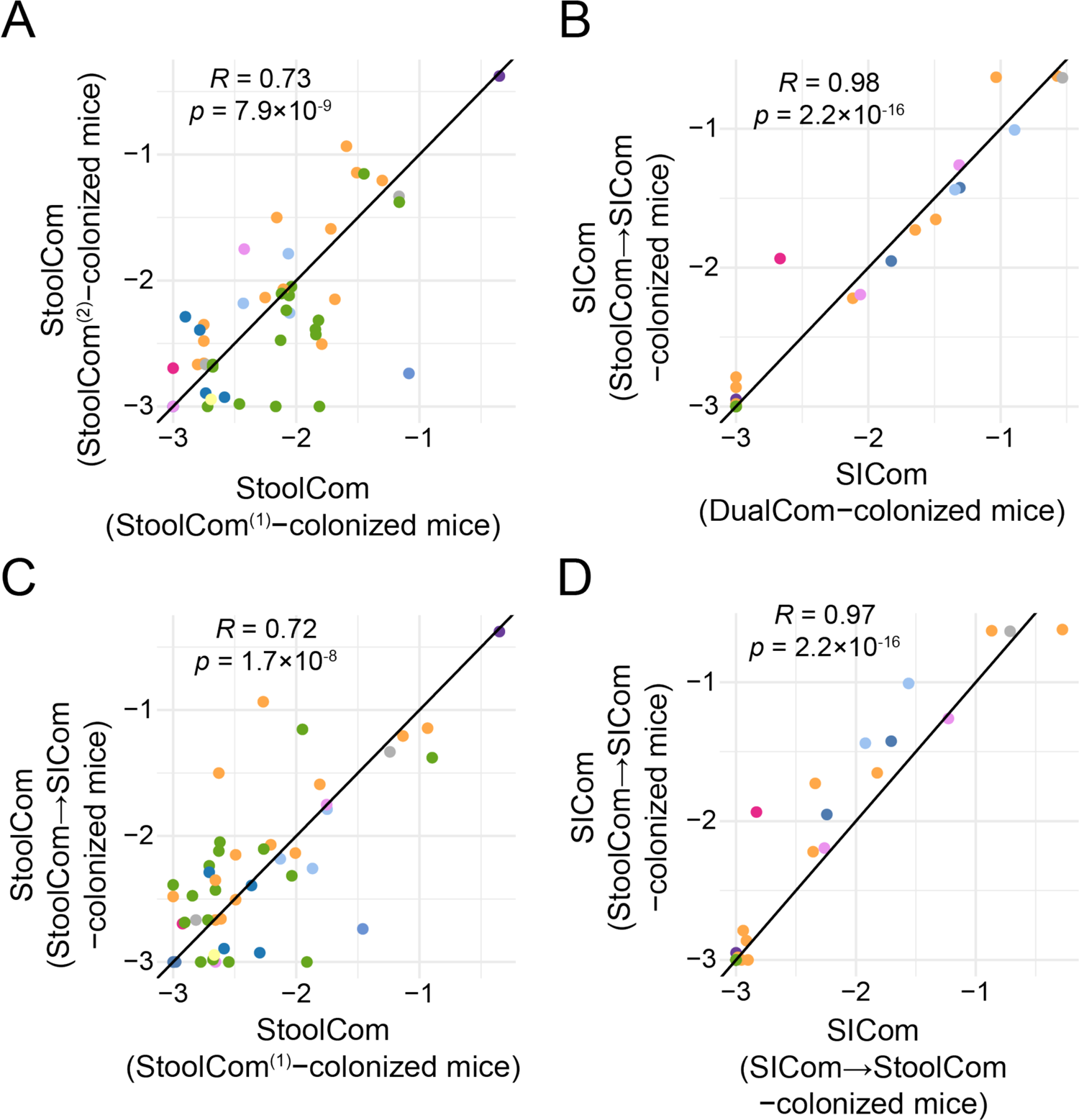
Inocula used for gavaging mice were similar across experiments. A,C) The StoolCom inoculum used to gavage StoolCom^(1)^-colonized mice (Fig. 2) was correlated at the ASV-level to the inocula used to colonize (A) StoolCom^(2)^-colonized mice (Fig. 3B,C) and (C) StoolCom→SICom-colonized mice (Fig. 3B,D). B,D) The SICom inoculum used to initially gavage StoolCom→SICom-colonized mice (Fig. 3B,D) was correlated at the ASV-level to the inocula used to (B) gavage DualCom^(1)^-colonized mice (Fig. 2) and (D) to cross-colonize SICom→StoolCom-colonized mice (Fig. 3B,D).

**Figure S4:**
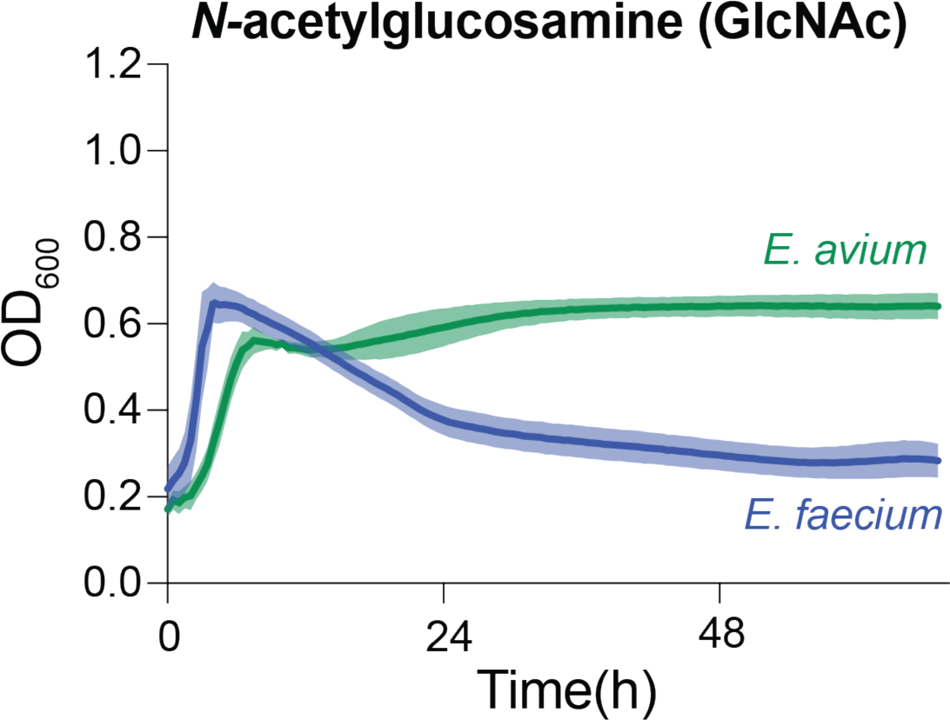
The *E. avium* isolate grew better than *E. faecium* on the mucin component N-acetylglucosamine (GlcNAc). *E. avium* and *E. faecium* reached a similar maximum optical density (OD), but *E. avium* maintained a high OD whereas *E. faecium* OD decreased dramatically. Shaded regions represent 1 standard error of the mean (*n*=3).

## Supplementary Tables

**Table S1: Isolates in the Small Intestinal Isolate Library.** Shown are the acquisition method, sample type (human/mouse stool/intestines or *in vitro* community), medium used for isolation, top BLAST annotation, and taxonomy.

**Table S2: Species likely to be enriched in the small intestine based on cross-colonization using SICom and StoolCom.** Shown in the log_2_(fold change) from the SI versus feces based on summed read counts across all mice. These data were used to inform the construction of synthetic communities.

**Table S3:** Synthetic community membership.

